# H_2_ dependent modulation of Mtr activity by small protein MtrR in *Methanosarcina mazei*

**DOI:** 10.1101/2025.05.30.657017

**Authors:** Tim Habenicht, Bjarne Hastedt, Liam Cassidy, Claudia Kießling, Andreas Tholey, Jan M. Schuller, Ruth A. Schmitz

**Author notes:** Corresponding author; Ruth A. Schmitz **Email**. **Author Contributions:** RAS initiated the project. TH and RAS designed the experiments. TH performed the majority of the experiments. BH and CK contributed to the cloning of expression mutants and *in vivo* interaction experiments. LC and AT performed LC-MS/MS analyses of MtrR interacting proteins. TH wrote the manuscript with RAS. TH, RAS, and JMS interpreted and finalized the results. RAS supervised the research and provided resources and funding. All authors approved the submitted version. We dedicate this work to Gerhard Gottschalk, a leading pioneer on methanoarchaea, on the occasion of his 90^th^ birthday with deep appreciation for his lifelong commitment and impact on the field.

## Abstract

Until recently, small open reading frame (sORF)-encoded proteins of fewer than 100 amino acids, have attracted increasing attention over the past decade after being overlooked due to limitations in conventional detection methodologies. While numerous previously unannotated sORFs have recently been identified in the mesophilic archaeal model system *Methanosarcina mazei*, the physiological roles of most of their encoded small proteins remain unknown. We report here the functional characterization of sORF16 encoded small protein MtrR (49 amino acids) and show that it localizes oligomerically at the cytoplasmic membrane. There, it interacts with and influences the activity of tetrahydrosarcinapterin S-methyltransferase (Mtr), a key membrane-bound complex involved in energy metabolism. *In vitro* interaction and *in vivo* copurification assays revealed interactions between MtrR and the Mtr-complex, and microscale thermophoresis showed specific interactions with the MtrA subunit. Mutant strains lacking sORF16 exhibited significantly impaired growth in the presence of molecular hydrogen (H_2_), irrespective of the carbon source. We posit that by modulating the activity of the Mtr-complex, MtrR enables the archaeon adapt to changing environmental H_2_ conditions.

## Introduction

Small proteins (< 100 amino acids [aa]) play important roles in prokaryotic cellular processes, *e*.*g*., metabolism, stress response, regulation of transcription and translation (Gray et al., 2022; Jordan et al., 2023; Storz et al., 2014; Weidenbach et al., 2022). Many small proteins, however, have been overlooked due to limitations in biochemical detection. As such, the small proteomes of most microorganisms, especially archaea, remain vastly understudied. Recently, small protein catalogs were generated via systematic genome-wide identification methods like ribosome footprinting (Ribo-Seq) and liquid chromatography-coupled tandem mass spectrometry (LC-MS/MS) (Cassidy et al., 2021; Hadjeras et al., 2023; Tufail et al., 2024). Functional analyses of most of these newly identified small proteins, however, remain scarce or lacking entirely (Gelsinger et al., 2020; Gray et al., 2022; Hadjeras et al., 2023; Jordan et al., 2023; Storz et al., 2014; Weidenbach et al., 2022).

The findings of studies employing mass spectrometry and Ribo-Seq analyses to identify novel small proteins encoded by small ORFs (sORFs) in the archaea *Haloferax volcanii* (Hadjeras et al., 2023) and *Methanosarcina mazei* Gö1 (Tufail et al., 2024) were recently reported. Forty-eight sORFs were identified in *H. volcanii*. These include HVO_2753 and HVO_0758 (Hadjeras et al., 2023; Üresin et al., 2023), which encode small zinc finger proteins (59 and 56 aa) that influence swarming activity and biofilm formation, respectively (Üresin et al., 2023; Zahn et al., 2021). Prior to the advent of LC-MS/MS, 69 small proteins had been identified in the methanoarchaeal model system of *M. mazei* (Cassidy et al., 2021; Prasse et al., 2015; Weidenbach et al., 2022), only a handful of which were functionally characterized. These include sP36 (61 aa), which regulates the ammonium transporter AmtB_1_ (Habenicht et al., 2023), and sP26 (23 aa), which modulates the activity of glutamine synthetase GlnA_1_ (Gutt et al., 2021). Ribo-Seq-coupled LC-MS/MS analyses unveiled 314 previously unannotated small ORFs, many of which were regulated according to nitrogen availability (Tufail et al., 2024).

Methanoarchaea contribute significantly to the global biotic production of methane, which is produced anoxically via acetoclastic, hydrogenotrophic, and/or methylotrophic pathways (Costa & Leigh, 2014). In acetoclastic methanogenesis, acetate is converted to methane and carbon dioxide. Here, cleavage of the C-C bond is catalyzed by the Acetyl-CoA decarbonylase/synthase complex (ACDS; (Grahame, 1991). In the hydrogenotrophic pathway, carbon dioxide is reduced to methane using molecular H_2_ as an electron donor. In a similar vein, the methylotrophic pathway exploits methylated compounds (*e*.*g*., methanol, methylamines, methyl sulfides) to reduce the methyl groups to CH_4_ (Costa & Leigh, 2014; Uwe Deppenmeier et al., 1996; Hippe et al., 1979; Huang et al., 2020; Mand & Metcalf, 2019). All three pathways rely on (i) the heterosulfide reductase (Hdr) to reduce the CoM-S-S-CoB heterodisulfate and thereby regenerate CoM and CoB, and (ii) final methane formation via methyl-coenzyme-M reductase (Hedderich et al., 2005; Jaun & Thauer, 2007).

Members of the genus *Methanosarcina* possess the unique ability to deploy all three pathways for methanogenesis (Hippe et al., 1979; Mand & Metcalf, 2019). As such, *Methanosarcina* species exhibit high levels of adaptability regarding available carbon and energy sources, in contrast to species of more specialized methanoarchaeal genera, *e*.*g*., *Methanobrevibacter* and *Methanococcus* (primarily hydrogenotrophic), *Methanosphaera* (methylotrophic), and *Methanosaeta* (acetoclastic) (Liu & Whitman, 2008). In *Methanosarcina*, these three pathways are regulated depending on environmental conditions, whereby energy output differs depending on the substrates available (Liu & Whitman, 2008). Methanogenesis using H_2_ in concert with CO_2_ (−136 kJ/mol CH_4_) or methylated substrates (−108 kJ/mol CH_4_) is far more exergonic than acetoclastic methanogenesis (−65 kJ/mol CH_4_; (Yi et al., 2023).

In the presence of H_2_, *M. mazei* grows hydrogenotrophically, reducing CO_2_ to methane (C_1_-metabolic pathway). This stepwise reduction of CO_2_ is initiated by its reduction to a formyl group bound to methanofuran (MFR). This formyl group is then transferred to tetrahydrosarcinapterin (H_4_SPT) and is further reduced to methenyl-H_4_SPT. Subsequent reduction yields methyl-H_4_SPT, the methyl group from which is then transferred to coenzyme M (CoM) by the membrane-bound tetrahydrosarcinapterin S-methyl transferase (Mtr)-complex. In the final step of methanogenesis, the methyl group is reduced to methane by methyl-CoM reductase. Electrons required for all reductions are supplied by H_2_ oxidation. In the absence of sufficient H_2_, *M. mazei* uses methylated compounds as carbon and energy sources (methylotrophy). In the absence of the electron donor H_2_, however, reducing equivalents must originate from an alternative source. In this case, a fraction of the methyl groups is oxidized to CO_2_, providing the electrons required for the reduction of further methyl groups to methane. This disproportionation process involves reversing part of the C_1_ metabolic pathway, where enzymes that normally reduce C_1_ compounds now function in reverse and oxidize methyl groups to CO_2_. This process commences with the transfer of the methyl group from methyl-CoM to H_4_SPT, catalyzed by the Mtr-complex operating in reverse (Uwe Deppenmeier et al., 1996).

Hitherto, our understanding of substrate-dependent regulation of metabolic pathways in response to available carbon and energy sources in *Methanosarcina* spp. has remained limited to the transcriptional level. In *M. acetivorans*, for example, expression of methyltransferase MtaC1 and MtaB1, which transfer the methyl group of methanol to CoM, are transcriptionally upregulated when methanol is present. When acetate is the sole carbon and energy source, however, expression of the ACDS complex is upregulated on the transcriptional level (Li et al., 2007). The molecular mechanisms underlying these regulatory processes at the transcriptional level often remain poorly understood. In addition, post-transcriptional and/or post-translational regulation has not been reported yet. Energy dependent anabolic processes might also influence the regulation of pathways in involved in energy metabolism, *e*.*g*., type of nitrogen source and availability (Yi et al., 2023).

We examined the physiological role of the 49 aa protein encoded by small ORF16, henceforth referred to as MtrR, which is transcribed and translated in significant amounts in *M. mazei* (Tufail et al., 2024). We show that, despite being devoid of transmembrane domains, small protein MtrR localizes exclusively at the cytoplasmic membrane, due to interaction with subunit A of the membrane-bound Mtr complex (MtrA). Based on biochemical and genetic analyses of this small protein, we posit that MtrR is responsible for H_2_-dependent modulation of Mtr complex activity, most likely by impeding the methyl transfer from CoM to H_4_SPT via direct protein-protein interactions.

## Results

### Localization of oligomeric MtrR on the cytoplasmic membrane

When MtrR was fused to a C-terminal his-tag (MtrR-His_6_), heterologously expressed in *Escherichia coli* and purified via Ni-NTA immobilized metal affinity chromatography (IMAC), elution fractions obtained at low imidazole concentrations yielded dimeric MtrR-His_6_ (∼ 14 kDa) and distinct proteins of greater molecular mass. The latter were likely larger oligomers that were not eluted at higher imidazole concentrations (Fig. 1A). Subsequently, size exclusion chromatography (SEC) yielded a purified fraction of homogeneous MtrR-His_6_ (Fig. 1B). Since the aa sequence of MtrR does not contain tryptophan, tyrosine, or cysteine residues, the SEC exhibits no visible peaks in the UV_280nm_ chromatogram. Consequently, exact determinations of molecular masses are difficult. Calibration with standard proteins suggested that the smallest MtrR-His_6_ fraction (13 ml) corresponds to a molecular mass of approximately 10 kDa, representing dimeric MtrR-His_6_ (supplemental Fig. S1). In preceding elution fractions (11 and 12 ml), additional larger oligomers of MtrR were detected (Fig. 1B). We conclude that MtrR forms dimers and larger oligomeric structures when in isolation.

**Figure 1.**
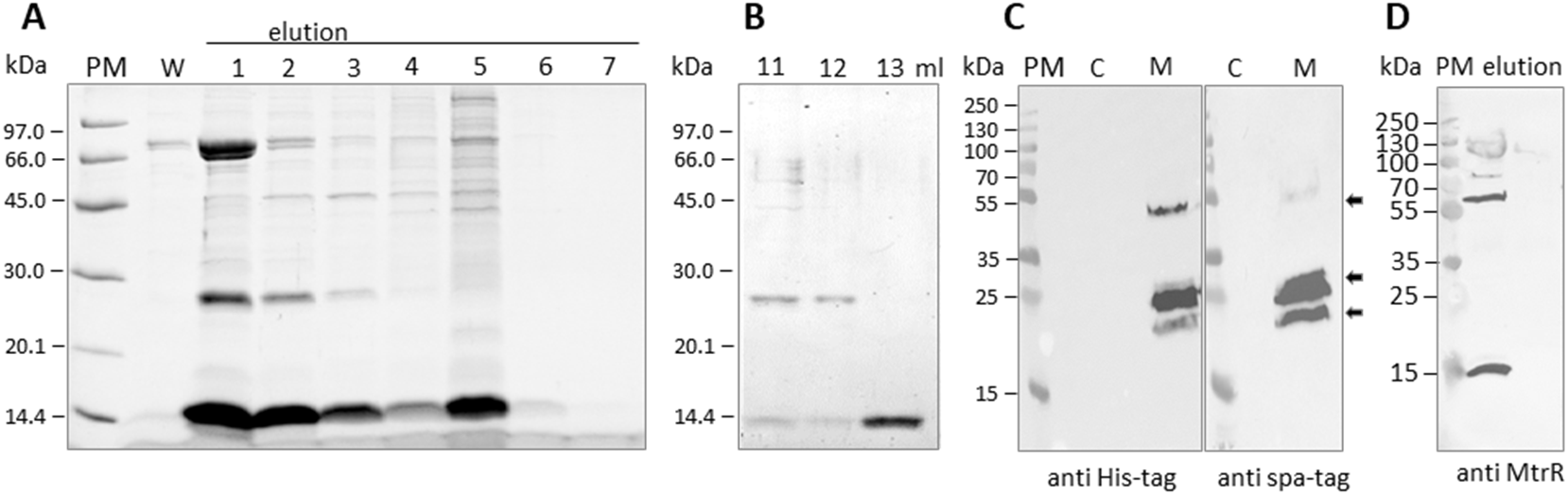
Purification and cellular localization of MtrR. **A:** SDS PAGE of IMAC purification of heterologously expressed MtrR-His_6_. W: final wash fraction; lanes 1-4: 100 mM imidazole; lanes 5-7, 250 mM imidazole. **B:** SEC purification of MtrR-His_6_ (Enrich SEC 70 column; BioRad, Hercules, CA USA) for downstream applications; a standard protein ladder (BioRad size-exclusion standard) was used for calibration, depicted are fractions 11 - 13 ml. The 13 ml elution fraction corresponds to a molecular mass of roughly 10 kDa (supp. Fig. S1). **C:** Exponentially growing *M. mazei* cells expressing MtrR-His_6_ or spa-MtrR from plasmids pRS2087 and pRS1693, respectively, were harvested, fractionated, and subcellular fractions analyzed by western blot using antibodies targeting either the His- or strep-tag. MtrR presence in the membrane fraction is denoted by arrows, corresponding to proteins (complexes) with a molecular mass of approximately 20, 25, and 50 kDa. C, cytoplasmic fraction; M, membrane fraction. The SDS-PAGE was loaded with 0.4 mg protein of both subcellular fractions. PM, PageRuler Plus Prestained Protein Ladder (Thermo Fisher Scientific, Waltham, MA USA). **D:** *In vivo* copurification: MtrR-His_6_ was constitutively expressed in *M. mazei* (pRS2087) and purified from the solubilized membrane fraction (2 % DDM) by IMAC. Elution fraction two of the purified MtrR complex was analyzed by western blot using antibodies generated against MtrR. Arrows denote MtrR detected in complexes of greater molecular mass (60, 80, and 130 kDa).

We analyzed the cellular localization and potential oligomerization of MtrR expressed from a plasmid in its native background. The gene encoding MtrR-His_6_ was cloned under the control of the constitutive promoter PmcrB, and sORF16 fused to an N-terminal spa-tag under the control of its native promoter. Cytoplasmic and membrane fractions of *M. mazei* cells were individually subjected to western blot analyses with antibodies directed against the respective tags. Regardless of the protein tag or promoter used, MtrR was detected exclusively in the membrane fraction and was absent in the cytoplasmic fraction (Fig. 1C). For both tagged versions, western blot signals corresponded to protein masses of approximately 20, 25, and 55 kDa, which likely represent stable homo-oligomeric forms of MtrR, and a 20 kDa protein degradation product. As MtrR is a hydrophilic, uncharged small protein that does not bear a transmembrane domain, the *in vivo* membrane localization of MtrR likely results from interactions with membrane-associated proteins.

MtrR-His_6_ was purified via IMAC from the solubilized membrane fraction (2% Dodecyl-β-D-maltosid (DDM)) of exponentially growing cells to confirm the oligomerization of MtrR expressed in *M. mazei* from a plasmid. Elution fractions analyzed by western blotting using specific antibodies generated against MtrR clearly showed the presence of MtrR in several larger (yet SDS-stable) oligomeric forms (roughly 60, 80, and 130 kDa) in addition to its native dimer (14 kDa, Fig. 1D).

### Small protein MtrR interacts with the tetrahydrosarcinapterin S-methyl transferase (Mtr) complex

The observation of MtrR localization at the membrane prompted the identification of target interaction proteins in the membrane. As such, we purified recombinant MtrR-His_6_ as bait for *in vitro* pull-downs of potentially interacting proteins from solubilized membrane fractions. Within three independent biological replicates, two protein signals (bands at 25 and 35 kDa) consistently co-purified with MtrR-His_6_, as shown in denaturing SDS-PAGE (Fig. 2 A). These additional bands were not detected when the membrane fraction was analyzed in the absence of MtrR-His_6_ (negative control; suppl. Fig. S2). These proteins were identified by mass spectrometry (with high confidence) as tetrahydrosarcinapterin S-methyl transferase (Mtr) subunits A and H in all three independent biological replicates (including controls; supp. Fig. S2). MtrA and MtrH are subunits of the large trimeric Mtr-complex (MtrABCDEFGH_2_). Transferring the methyl group from methyl-H_4_SPT to the final cofactor, *i*.*e*. CoM, Mtr is a key membrane-bound enzyme complex in energy metabolism. Of its eight subunits, MtrH and MtrA are the two most hydrophilic, the latter being completely soluble and capable of forming dimers linked to the membrane via interactions with subunits MtrB and MtrF (Reif-Trauttmansdorff et al., 2025). MtrA bears a large cytoplasmic domain and a small transmembrane helix, which anchors it securely to the stem of the trimeric Mtr complex.

**Figure 2.**
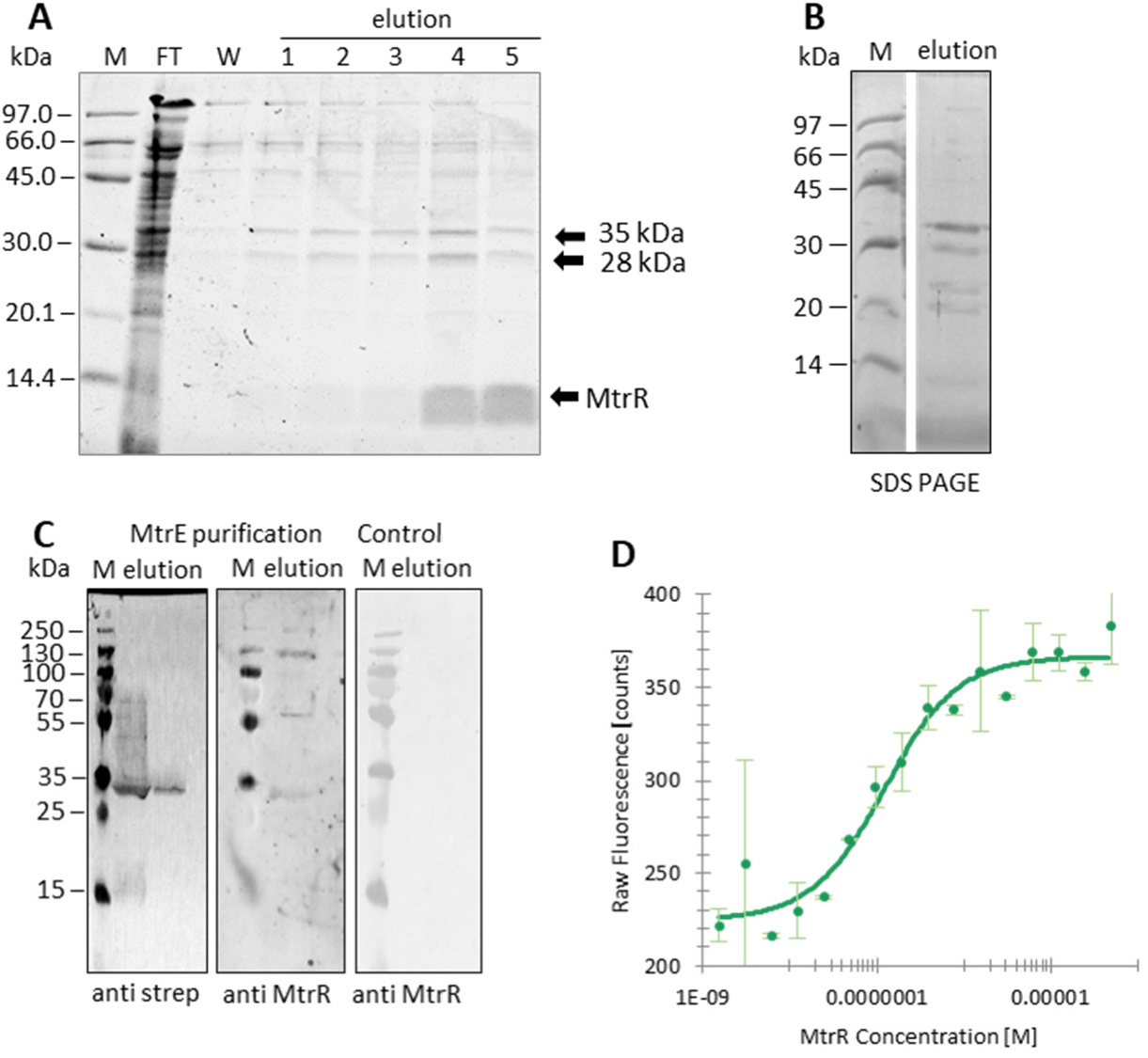
**MtrR interacts with the Mtr-complex, particularly MtrA. A: SDS-PAGE resulting from** pull-down analyses using purified MtrR-His_6_ as bait against solubilized *M. mazei* membrane fractions (2 % DDM). After 30 min incubation at RT, MtrR-His_6_ was purified by IMAC revealing two additional distinct protein bands, as indicated. **B and C:** *In vivo* co-purification of the Mtr complex from *M. mazei* using plasmid encoded strep-MtrE. **B**: Elutions of the strep-tactin purified complex were analyzed by SDS PAGE. Visible are the different copurified subunits of the Mtr complex **C:** Elutions of the strep-tactin purified complex were analyzed via western blot using antibodies directed against the strep-tag or MtrR. Results clearly show coelution of MtrR with the Mtr complex purified by strep-MtrE at approximately 30, 60, and 130 kDa. *M. mazei* WT cells not encoding any strep tagged proteins were used as a control (right panel). **D: Affinity analysis of the MtrR-MtrA interaction by microscale thermophoresis**. 20 nM RED labeled heterologously expressed strep-MtrA was incubated with His-MtrR in concentrations ranging from 3 to 100 μM, calculated based on monomeric molecular mass. Based on two biological replicates, K_D_ was estimated to be 110 ± 30 nM.

To verify interactions with the Mtr complex, a reverse approach was tested via *in vivo* co-purification using strep-MtrE constitutively expressed from a plasmid (pRS1743) in *M. mazei* to purify the Mtr complex and potential interaction partners, such as MtrR from solubilized cell extract (1% LMNG; (Reif-Trauttmansdorff et al., 2025). The elution fraction resulting from the purified strep-tagged MtrE analyzed on SDS PAGE yielded the correct protein pattern (as reported in (Reif-Trauttmansdorff et al., 2025)), including the 28 and 35-kDa subunits MtrA and MtrH, respectively (Fig. 2B). When this elution fraction was subjected to western blot analysis employing antibodies against strep-tag and MtrR, MtrE emerged primarily as a monomer (30 kDa), and also in 50, 60, and 70 kDa complexes. These likely correspond to complexes in which different Mtr subunits interact (Fig. 2C, left panel). When using specific antibodies, MtrR was clearly detected in the purified Mtr-complex (Fig. 2C, mid panel), in stark contrast to its respective control (Fig. 2C, right panel). MtrR was present as a dimer (∼ 14 kDa), as well as associated with complexes of 30, 60, and 130 kDa. This pattern closely mirrors the one observed for MtrR-His_6_ purification (Fig. 1D), showing a consistent oligomerization pattern of MtrR in *M. mazei*. Coelution confirms that MtrR copurifies with the Mtr complex, as corroborated by pull-down analyses (Fig. 2A), and hints at the presence of larger MtrR oligomers in the Mtr complex.

The soluble domain of the MtrA subunit (M1 – S168) without the hydrophobic membrane anchor (E169 - R240) was cloned as a strep-tagged version in *E. coli* and purified by affinity chromatography. Interaction analysis with purified, fluorescently labeled MtrA (MtrA-RED, 20 nM) was performed via microscale thermophoresis (MST) using unlabeled MtrR Affinity analyses conducted using MO.Affinity (Nanotemper, Munich, Germany) yielded an estimated dissociation constant of *K*_D_ = 110 ± 30 nM. In each of two independent biological replicates, MST analyses detected strong interactions between MtrA-RED and MtrR *in vitro*, supporting the results of *in vitro* pull-down and *in vivo* copurification experiments.

### MtrR modulates activity of the Mtr complex in vivo

We devised a genetic approach to garner insights into potential regulatory impacts of MtrR on the Mtr-complex activity upon binding. Overproduction of MtrR-His_6_ in *M. mazei* from a plasmid (pRS2087) did not affect growth under standard conditions with methanol as the sole carbon and energy source. In addition, a chromosomal deletion mutant of small ORF16 was constructed via allelic replacement through homologous recombination, replacing small ORF16 with a puromycin resistance cassette (*pac*). The successful generation of this mutant strain (*M. mazei* Δ*sORF16*) was verified by southern blot analysis (supp. Fig. S3), and its growth characteristics under different conditions - varying the availability of carbon and energy sources as well as of molecular hydrogen (H_2_) - was evaluated and compared to the wild type strain (Fig. 3). No effect was observed under standard growth conditions with methanol as the sole carbon and energy source with a gas phase consisting of N_2_/CO_2_ (v/v 80/20) (Fig. 3A). Under those conditions (favoring methylotrophic methanogenesis), roughly 25% of the available methanol is oxidized to CO_2_ to provide electrons to reduce the remaining 75% of the methanol fraction. Upon altering the N_2_/CO_2_ gas phase to 100% H_2_, the wild type grew faster and to a greater final cell density (Fig. 3A and B, blue lines). This improved growth performance is a consequence of the sole use of molecular hydrogen as the electron donor, consequently reducing methanol exclusively to methane. In contrast, the *M. mazei* Δ*sORF16* mutant strain does not capitalize on the presence of H_2_ in the gas phase, exhibiting identical growth behavior as that observed in the absence of H_2_. As might be expected, this resulted slower growth and lower cell density than the wild type strain (Fig. 3B, orange line). In addition, when growing in the absence of methanol with CO_2_ and H_2_ as the main carbon and energy sources in the gas phase (favoring hydrogenotrophic methanogenesis), the *M. mazei* Δ*sORF16* mutant strain once again exhibited slower growth and lower final cell density than its wild type counterpart (Fig. 3C, orange line). The results of both genetic and biochemical experiments strongly suggest a regulatory role for small protein MtrR in energy metabolism at the methyl transfer step catalyzed by the Mtr complex in the presence of molecular H_2_.

**Figure 3.**
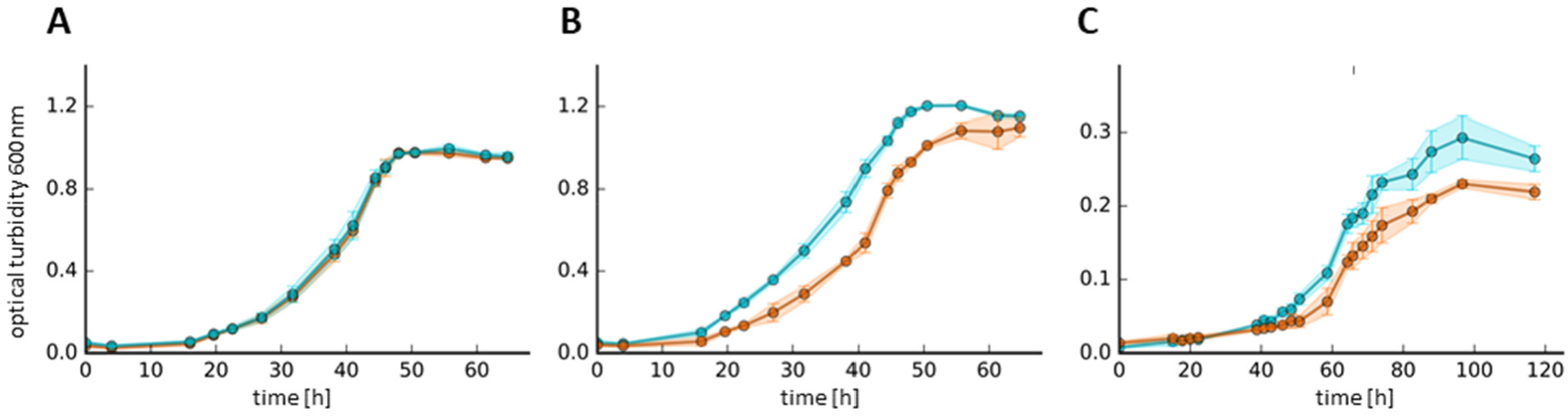
Growth of the *sORF16* deletion strain under different conditions. The growth behavior of *M. mazei* sORF16 chromosomal deletion mutant was studied under different growth conditions. Cultures (50 ml) were inoculated and incubated in closed 80 ml serum bottles under the respective gas phase. All growth curves depict the mean of three biological replicates each with several technical replicates, standard deviation is indicated as error bars. Proliferation of wt *M. mazei* is charted in blue and the deletion mutant ΔsORF16 in orange. **A:** Growth on 150 mM MeOH with a N_2_/CO_2_ (vol/vol, 80/20) gas phase. Inoculated from precultures grown on 150 mM MeOH with a N_2_/CO_2_ (vol/vol, 80/20) gas phase (methylotrophic growth). **B:** Growth on MeOH (150 mM) with a H_2_ gas phase (100%). Inoculated from precultures grown on 150 mM MeOH with a N_2_/CO_2_ (vol/vol, 80/20) gas phase. **C:** Growth on H_2_/CO_2_ (vol/vol, 80/20) as sole energy and carbon source in the gas phase (favoring hydrogenotrophic methanogenesis). Inoculated from precultures grown without MeOH with a H_2_/CO_2_ (vol/vol, 80/20) gas phase. Please note the different scale of the y-axis in **C**.

## Discussion

Aiming to functionally characterize the 49-aa small protein MtrR encoded by small ORF16, which was identified in *M. mazei* through ribosome profiling and mass spectroscopy (Tufail et al., 2024), we studied the small protein by biochemical and genetic approaches. We obtained strong experimental indications that the 5.5 kDa small protein regulates the membrane-bound Mtr complex. This work demonstrates that MtrR, which lacks a membrane domain itself, is interacting with the membrane-bound Mtr complex resulting in the localization of oligomeric MtrR at the cytoplasmic membrane (Fig. 1C). In pull-down experiments using purified MtrR-His_6_ and solubilized membranes, native MtrA and MtrH were copurified and identified by LC-MS/MS analysis (Fig. 2A). MtrR interaction with the Mtr complex was further confirmed by native and reversed copurification approaches (Fig. 2 BC) as well as by MST analysis between MtrR and MtrA showing high binding affinity (Fig. 2D).

The large, membrane-associated, multi-enzyme Mtr complex is crucial to methanogenesis. In many methanogenic archaea, it facilitates the exergonic transfer of the methyl group from methyl tetrahydromethanopterin (CH_3_-H_4_MPT) to coenzyme M (HS-CoM; ΔG = -30 kJ/mol;(Becher & Muller, 1994; Mand & Metcalf, 2019). In *M. mazei*, instead of being transferred from methanopterin, the methyl group is transferred from H_4_SPT, which is a derivative of methanopterin with a conjugated L-glutamic acid (Van Beelen et al., 1984). This Mtr-mediated reaction is coupled to sodium ion translocation over the cytoplasmic membrane, furthering energy conservation via establishment of an ion gradient across the cytoplasmic membrane (Perski et al., 1982). The large trimeric Mtr-complex consists of eight different subunits, MtrABCDEFGH_2_ (Aziz et al., 2024). MtrB, C, D, E, F, and G are transmembrane proteins bearing relatively small extracellular and/or cytoplasmic moieties. MtrA consists of a transmembrane helix bound to the membrane via the MtrBCDEFG complex, and another approximately 18-kDa soluble domain. While MtrH is a soluble protein, is associates with the membrane in dimeric form via interactions with MtrB and MtrF, which comprise the central transmembrane stalk (suppl. Fig. S4F). Bearing the binding site for CH_3_-H_4_SPT, MtrH transfers the methyl group to a MtrA-bound cobalamin. Following the methyl transfer, the methyl-cobalamin-carrying MtrA dissociates from MtrH and binds the MtrCDE subcomplex, which houses cofactor HS-CoM, onto which the methyl group is transferred (suppl. Fig. S4F). MtrA is predicted to behave as a shuttle for methyl groups, tirelessly switching back-and-forth between MtrH and the membrane-bound MtrCDE (Aziz et al., 2024; Uwe Deppenmeier et al., 1996; Gottschalk & Thauer, 2001; Reif-Trauttmansdorff et al., 2025).

In *Methanosarcina*, the Mtr-complex can catalyze methyl transfer in both directions depending on environmental conditions, thereby facilitating methylotrophic, hydrogenotrophic, and/or acetoclastic methanogenesis. When growing on methanol in the absence of H_2_ as an electron donor, *M. mazei* disproportionates the methanol, where one mol of methanol is oxidized to generate the electrons required for the reduction of three other mol of methanol (Fig. 4C). The first step of this oxidation is the transfer of the methyl group from CH_3_-CoM to H_4_SPT, performed by the Mtr-complex. This is simply the Mtr-mediated reductive reaction described above run in reverse. This reversed reaction is thermodynamically unfavorable (ΔG = +30 kJ/mol), and as such needs to be driven by electrochemical membrane potential. One can speculate that the direction of this reaction is governed by environmental conditions, such as carbon sources and H_2_ availability. However, a molecular mechanism underlying this regulation has yet to be reported.

**Figure 4.**
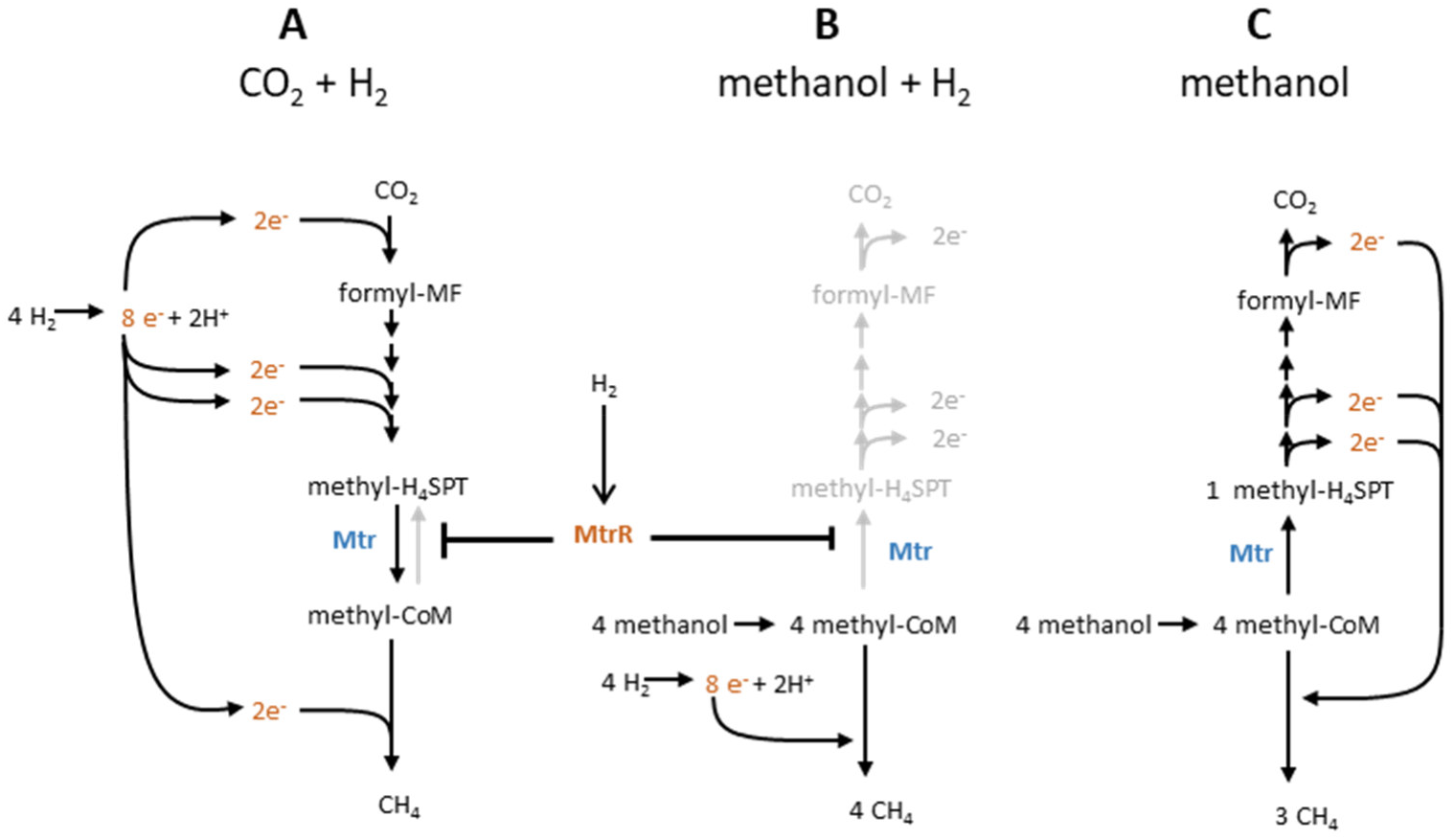
Hypothetical model of Mtr regulation by MtrR. Simplified schematic cartoons of methanogenesis in *M. mazei* under three different conditions: **A**: CO_2_ as sole carbon and energy source in the presence of molecular H_2_. **B/C**: Growth on methanol with H_2_ (**B**) and without H_2_ (**C**) in the gas phase. We posit that MtrR and H_2_ are required to drive methyl transfer from CH_3_-H_4_SPT to CoM (*i*.*e*., in a reducing direction), likely via protein-protein interactions.

As MtrR interacts with the Mtr-complex in *M. mazei* and is conserved exclusively in the *Methanosarcina* genus, which can capitalize a variety of carbon and energy sources, we posit a regulatory function for MtrR in energy metabolism ultimately governed by the availability of various carbon sources and electron donors. Consistent with computational predictions obtained using AlphaFold3, which provide tentative structural insights into potential interactions between MtrR and the Mtr complex (suppl. Fig. S4), MST experiments demonstrated that MtrR binds the MtrA subunit specifically. MtrR (aa 24 - 28) might interact with the MtrA surface opposite the cobalamin binding site. Here, a flexible loop tethers MtrA’s catalytic domain to its membrane anchor (see suppl. Fig. S4). While these interactions may not hinder transfer of the methyl group in general, they might cause MtrA to favor transfer of the methyl group to CoM instead of H_4_SPT.

This hypothesis, where MtrR invokes the favoring the methyl transfer from CH_3_-H_4_SPT to CoM, is in agreement with the growth characteristics of the chromosomal deletion mutant (Δ*sORF16*). Using methanol as both a carbon and energy source, wild type *M. mazei* grows better in the presence of H_2_, as it readily provides the electrons needed to reduce the methanol to methane (Fig. 3). As such, no disproportionation of methanol is required (Fig. 4C) and all methanol is reduced to methane (see Fig. 4B). However, the Δ*sORF16* mutant strain fails to capitalize fully on these readily available electrons, resulting in the in continued partial oxidation of methanol, less energy conservation, and slower growth (Fig. 3B). When carbon dioxide and molecular hydrogen are the sole sources of carbon and energy (thus favoring hydrogenotrophic methanogenesis), the Δ*sORF16* mutant strain exhibits much slower growth its wild-type counterpart (Fig. 3C). This suggests that in the absence of MtrR, the Mtr complex partially catalyzes the transfer of the methyl group from CoM to H_4_SPT as part of the methyl-oxidizing pathway, resulting in lower energy yields and slower growth.

We conclude that in the presence of H_2_, small protein MtrR plays an important role in the post-translational regulation of the membrane-bound Mtr complex, a key driver of methanogenesis. MtrR causes the Mtr complex to favor methyl transfer from CH_3_-H_4_SPT to CoM (reducing direction), most likely via protein-protein interactions. This unique post-translational regulation of the Mtr complex by a small protein renders superior fitness upon the organism in the presence of H_2_, regardless of whether methanol or carbon dioxide is available as a carbon source (Fig. 4). Moreover, the findings argue for a direct link between small protein modulation, energy conservation, and environmental sensing in methanoarchaea. We speculate that, based on H_2_ presence and/or cellular energy demand, MtrR is upregulated or activated and subsequently regulates the Mtr complex by binding to MtrA. Future investigations into this regulation at the molecular and structural level, as well as the signaling pathway for H_2_ availability, will bolster our understanding of the important regulation of the key enzyme Mtr. Acquired detailed knowledge on this regulation might in the long term allow managing production of methane as a climate friendly alternative to fossil fuels.

## Methods

### Protein structure prediction

Protein structures were predicted using AlphaFold3 via the AlphaFold3 Server (Abramson et al., 2024; Jumper et al., 2021).

### Construction of strains and plasmids

#### Small ORF16 genomic deletion

The 1-kb genomic regions flanking the small ORF16 gene on both sides were amplified from *M. mazei* Gö1 genomic DNA using specific primers (Eurofins, Ebersberg, Germany) listed in Table S2. A puromycin resistance cassette (PurR; (Metcalf et al., 1997), encoding the puromycin *N*-acetyltransferase gene from *Streptomyces alboniger* coupled to *M. mazei* constitutive promoter pmcrB, was excised from pRS207 (Table S1) with *Bam*HI. The 1-kb upstream and downstream fragments were inserted into plasmid pMCL210 with restriction enzymes *Kpn*I and *Not*I. Plasmid pMCL210 (Nakano et al., 1995) was used as a suicide vector for allelic replacement in *M. mazei* (Hüttermann & Schmitz, 2024). The resulting plasmid, designated pRS2037, was linearized using *Sal*I and transformed into *M. mazei* via liposome-mediated transformation (Ehlers et al., 2005). The small ORF16 gene was replaced with the *pac*-cassette using the 1kb flanking regions for double homologous recombination. Successfully recombined cells bearing the small ORF16 deletion were selected in puromycin-containing medium and plated to obtain single mutant clones (Uwe Deppenmeier et al., 2002). Success of the deletion was confirmed via southern blotting using specific probes against the small ORF16 gene and the *pac* cassette (Fig S3).

#### Small ORF16 expression mutant

The gene encoding MtrR was amplified from *M. mazei* genomic DNA using specific primers (Eurofins, Ebersberg, Germany; Table S2) and subjected to restriction ligation cloning into pET21a with *Nde*I and *Not*I restriction enzymes (New England Biolabs, Ipswich, MA, USA) and T4 ligase (Thermo Fisher Scientific, Waltham, MA, USA). The resulting plasmid, *i*.*e*. pRS1834, was transformed into competent (Inoue et al., 1990) *E. coli* BL21 pRIL (Table S1) for heterologous expression. The small ORF16-His_6_ fusion was amplified from pRS1834 and cloned into pRS1807 downstream of *M. mazei* constitutive promotor pmcrB with *Bam*HI and *Not*I (New England Biolabs, Ipswich, MA, USA) and T4 ligase (Thermo Fisher Scientific, Waltham, MA, USA). The resulting plasmid, deemed pRS2087, was then transformed into *M. mazei* via liposome-mediated transformation (Ehlers et al., 2005).

The nucleotide sequence of the soluble part (aa 1-168) of the *mtrA* gene without the transmembrane domain (aa 169-240) was amplified from *M. mazei* genomic DNA using specific primers (Eurofins, Ebersberg, Germany; Table S2) and cloned into pRS1736 with B*sa*I restriction enzyme (New England Biolabs, Ipswich, MA, USA) and T4 ligase (Thermo Fisher Scientific, Waltham, MA, USA). The resulting plasmid, named pRS2190, was transformed into competent (Inoue et al., 1990) *E. coli* BL21 pRIL (Table S1) for heterologous expression.

### Purification of heterologously expressed proteins

*E. coli* BL21 (DE3)/pRIL cultures containing the respective expression plasmid were grown in Terrific Broth (TB) medium under continuous shaking at 37 °C. At an optical turbidity of T600 = 0.8, protein expression was induced by adding IPTG to a final concentration of 100 μM. Following an additional incubation of 2 h, cells were harvested by centrifugation at 4,000 x g at 4 °C for 20 min. Cells were then resuspended in 6 mL phosphate buffer A (50 mM sodium phosphate, 300 mM NaCl, pH 8.0) and lysed by passing the French pressure cell two times with 80 N/mm^2^. To remove unlysed cells and cellular debris, the extract was centrifuged for an additional 30 min at 13,000xg at 4°C. His tagged proteins were purified via IMAC using Ni-NTA on gravity columns with 1 ml bed volume. The elution was conducted using phosphate buffer (50 mM sodium phosphate, 300 mM NaCl, pH 8.0) with imidazole in steps with 100 mM, 250 mM and 500 mM in phosphate buffer A. Strep-tagged proteins were purified with strep tactin sepharose (IBA Lifesciences, Göttingen, Germany) and eluted with 2.5 mM desthiobiotin in Tris/HCl buffer (100 mM Tris-HCl, 150 mM NaCl, 2.5 mM EDTA pH 8.0). The buffer was subsequently exchanged to phosphate buffer (50 mM sodium phosphate, 300 mM NaCl, pH 8.0). When necessary, additional purification was achieved via size exclusion chromatography (SEC) using a gel filtration column (Enrich TM SEC 650; BioRad, Hercules, CA, USA). Proteins were eluted at a flow rate of 1 ml/min in phosphate buffer (50 mM sodium phosphate, 300 mM NaCl, pH 8.0). MtrR-His_6_ was purified to homogeneity (Fig. 1B, 13 ml) and used to generate polyclonal antibodies (Davids Biotechnologie; Regensburg, Germany).

### Growth of *M. mazei* and subcellular fractionation

*M. mazei* was cultivated under standard conditions in either 1 L anaerobic minimal medium in 2 L sealed bottles or in 50 mL anaerobic minimal medium (U. Deppenmeier et al., 1990) in 80 mL serum bottles with a gas phase consisting of N_2_ and CO_2_ in an 80/20 (vol/vol,) ratio (2 bar). Immediately prior to inoculation, the medium was augmented with 150 mM methanol as a carbon source. For growth experiments, the gas phase was changed to H_2_/CO_2_ in an 80/20 (vol/vol) ratio or 100% H_2_ (2 bar) ± methanol. *M. mazei* cells were then harvested via centrifugation at 6,000 x g at 4 °C for 45 min, and cells were lysed using a dismembrator (Sartorius, Göttingen, Germany) at 1,600 rpm for 3 min in Tris buffer (50mM, pH 7.6). Subcellular fractionation was achieved via centrifugation at 200,000 x g at 4 °C for 1 h. The supernatant served as the cytoplasmic fraction. The pellet was washed in Tris buffer, re-centrifuged (in the same manner), solubilized by adding a final concentration of either 2% DDM or 1% LMNG and shaking for 1 h at 4 °C, and subsequently centrifuged again at 210,000 x g and 4 °C for 1 h. The resulting pellet served as the membrane fraction. To preclude the aggregation of membrane proteins, subcellular samples were not heated prior to SDS-PAGE loading.

### Pull-down from *M. mazei* and LC-MS/MS analysis

Heterologously expressed and purified MtrR-His_6_ was used as bait protein for *in vitro* pull-down analyses of potentially interacting proteins from solubilized membrane fractions. His_6_-MtrR was purified via IMAC and subsequent SEC to obtain protein fractions of high purity to be used as bait (see Fig. 1B). Membrane fractions of *M. mazei* were prepared by subcellular fractionation from exponentially growing cells at an optical turbidity of 0.6, as described above. The resulting membrane fraction (total protein content of 5 mg) was incubated with the bait protein (0.5 mg purified MtrR-His_6_), followed by IMAC purification of MtrR-His_6_ using 0.1 ml Ni-NTA agarose resin (Qiagen, Hilden, Germany). A membrane fraction (total protein content of 5 mg) without bait MtrR-His_6_ was treated in the same manner and served as a negative control. Potential interaction partners for small protein MtrR were identified from the elution via GeLC-MS. Briefly, triplicate samples and respective controls were separated across 12% SDS-PAGE and stained with Coomassie brilliant blue. To prevent aggregation of the membrane proteins, samples of membrane proteins were not heated prior to loading into the SDS-PAGE gels.

Protein bands present in the sample but absent in the control gels were excised, and comparable regions from the control gels were also processed. The excised fractions were cut into cubes *ca*. 1mm^3^, and de-stained via successive washes of acetonitrile and 100 mM ammonium bicarbonate buffer, pH 7.4. In gel reduction (10 mM dithiothreitol, 56°C, 1 hr) and alkylation (50 mM chloroacetamide, 20°C, 30 min) of the samples was performed, followed by overnight enzymatic digestion with sequencing-grade trypsin (20 ng/band, in 100 mM ammonium bicarbonate buffer, pH 7.4, 37°C). Following digestion, peptides in the aqueous solution were collected and two subsequent extractions were performed on the hydrophobic peptides: one with 60% acetonitrile + 0.1% trifluoroacetic acid and another in 100% acetonitrile. The peptides resulting from both extractions were then pooled with those resulting from the aqueous solution of the corresponding sample, priorto being dried via vacuum centrifugation.

Purified peptides were re-suspended in 3% acetonitrile + 0.05% trifluoroacetic acid immediately prior to LC-MS analysis. One-dimensional liquid chromatography separation of the peptides was performed on a Dionex U3000RSLC nanoHPLC (Thermo Fisher Scientific, Germany) across an Acclaim Pepmap100 C18 column (2μm particle size, id75 μm × 500 mm) coupled online to a Q Exactive Plus mass spectrometer (Thermo Fisher Scientific, Germany). Eluent A consisted of 0.05% formic acid while eluent B consisted of 80% acetonitrile + 0.04% formic acid. The separation continued over a programmed 60 min run. Initial chromatographic conditions were 4% eluent B in eluent A, for 3 min followed by linear gradients from 4% to 50% eluent B over 30 min, then 50 to 90% over 1 min, and 10 min at 90% eluent B. Following this, an inter-run equilibration of the column was conducted for 15 min at 4% eluent B. A 300 nl/min flow rate and 5 μl were injected per run.

Data acquisition was conducted in a data-dependent manner with full scan MS acquisition (300-1500 m/z, resolution 70,000) and subsequent MS/MS of the top 10 most intense ions via HCD activation at NCE 27 (resolution 17,500); dynamic exclusion was enabled (20 sec duration). Data analysis was performed using Proteome Discoverer (Ver 3.1.1.93) and the SequestHT search algorithm. MS raw files were searched against the *M. mazei* Gö1 database including all predicted SEP protein sequences (accession date 2024.06.19) and the cRAP list of common laboratory contaminants. Searches were performed with trypsin specificity and a maximum of two missed cleavages, methionine oxidation was set as a variable modification, and cysteine carbamidomethylation was set as a fixed modification. Raw data files can be found in the ProteomeXchange repository (identifier PXD064332 (Röst et al., 2014).

### Microscale thermophoresis

Proteins were purified to homogeneity by affinity chromatography using Ni-NTA agarose (Qiagen, Hilden, Germany) and labeled with the RED-NHS, 2nd generation, 650 nm fluorescent dye using the Monolith NT RED-NHS lysine labeling kit, according to the manufacturer’s protocol (NanoTemper, Munich, Germany). RED-labeled strep-tagged MtrA at 20 nM and MtrR-His6 in 16 different concentrations ranging from 3 μM to 100 nM (based on monomeric molecular mass) were used. Protein interactions were measured in standard capillaries (Nanotemper, Munich, Germany) at 100% excitation power and medium MST power (IR-laser intensity). Each interaction was tested in two biological replicates. MST traces were analyzed in MO. Affinity Analysis (Nanotemper, Munich, Germany).

## Acknowledgments

We thank the groups of AG Schmitz-Streit and AG Schuller for fruitful discussions and critical comments. This work was supported by the German Research Foundation (DFG) priority program SPP2002 “Small Proteins in Prokaryotes, an Unexplored World” RSCHM1052/20-1 and 20-2, RSCHM1052/19-2 to RAS; TH872/10-1 and 10-2 to A.T. J.M.S. acknowledges the DFG for an Emmy Noether grant (SCHU 3364/1-1), RTG 2937 and the European Union’s Horizon 2020 research and innovation programme (Two-CO2-One; grant agreement no. 101075992). The views and opinions expressed are those of the authors only and do not necessarily reflect those of the European Union or the European Research Council. Neither the European Union nor the granting authority can be held responsible for them.

