## Supplemental Figures and Tables for "H_2_ dependent modulation of Mtr activity by small protein MtrR in *Methanosarcina mazei*" (from another version of this preprint: version 1, 2025-05-30)

### Supplement

Content:

Suppl. Fig. S1: Size exclusion of MtrR and the size exclusion standard proteins.

Suppl. Fig. S2: Pull-down analysis using purified MtrR-$His_6$ as bait against solubilized *M. mazei* membrane fractions

Suppl. Fig. S3: Southern blot of the *M. mazei* WT and the Δ*sORF16* strain

Suppl. Fig. S4: Computational model of the $MtrABCDEFGH_2$-complex

**Suppl. Table S1: Used Strains and plasmids**

Suppl. Table S2: Used oligonucleotides

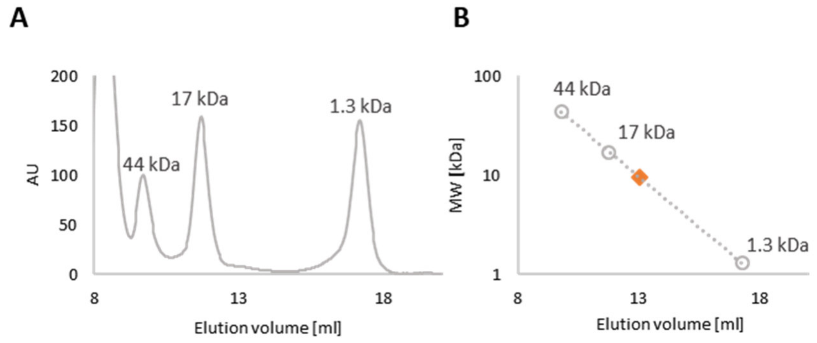


**Fig. S1: Size exclusion of MtrR and the size exclusion standard proteins**. **A:** A standard protein mix (Bio-Rad size exclusion standard; Bio-Rad, Hercules, USA) was used as size exclusion standard. The standard and the protein samples were run (Enrich 70, BioRad) with a flow rate of 1 ml $min^{-1}$ with 50 mM phosphate buffer (150 mM NaCl, pH 8.0). **B:** The three protein peaks which show a logarithmic relation are shown with the respective linear regression line. MtrR, which elutes at approximately 13 ml is indicated on the linear regression as an orange diamond (corresponding to approx. 10 kDa).

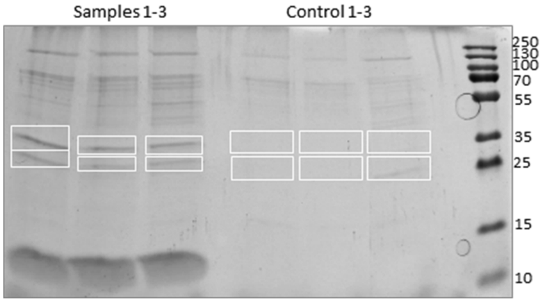


**Fig. S2: Pull-down analysis using purified MtrR-His$_6$ as bait against solubilized *M. mazei* membrane fractions (2% DDM)**. After 30 min incubation at RT, MtrR-His$_6$ was purified by IMAC. Two co-purifying proteins per (independent) biological sample were cut from the SDS gel as indicated by the white boxes. As negative control, solubilized *M. mazei* membrane fractions without addition of MtrR-His$_6$ were treated equally and corresponding parts of the gel were cut from the SDS PAGE as indicated. Both, samples and control gel parts were used for LC-MS/MS analysis.

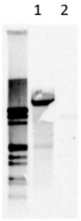


**Fig. S3: Southern Blot of the *M. mazei* WT (1) and the ∆*ORF16* strain (2)** with specific probes targeting the small ORF16. The signal is only present in the WT confirming the successful genomic deletion of sORF16.

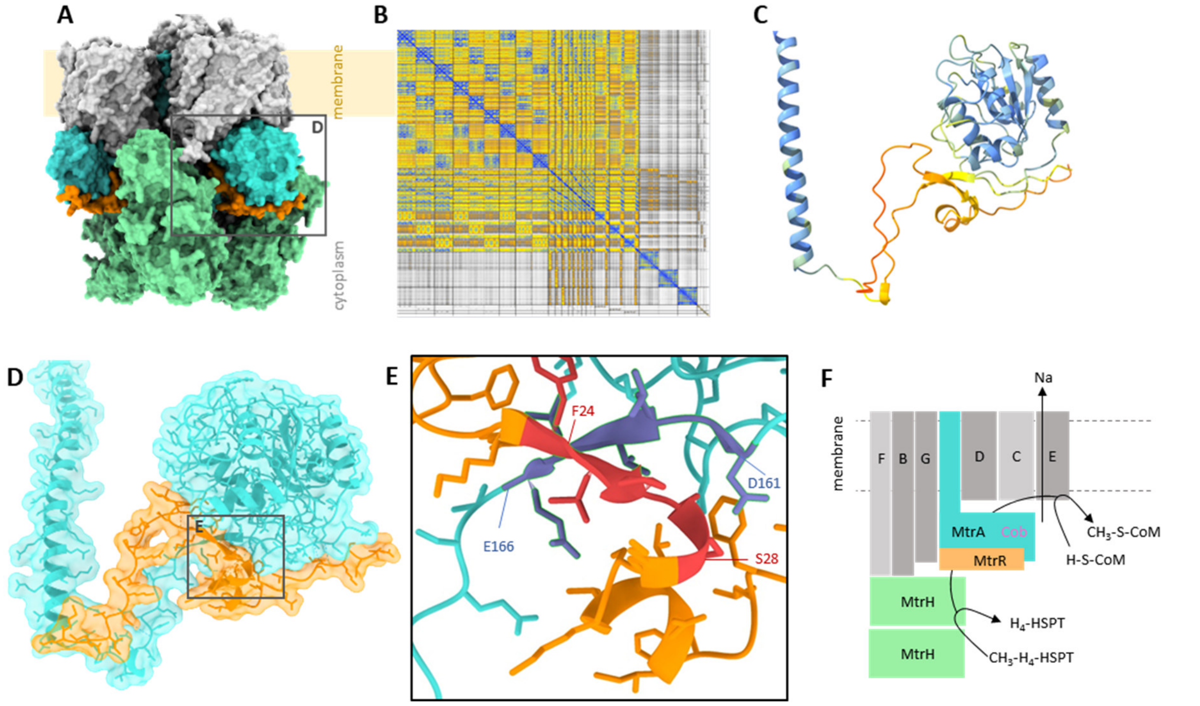


**Fig. S4: Computational model of the MtrABCDEFGH$_2$-complex** including MtrR predicted by AlphaFold3, analyzed using ChimeraX and Alphabridge. Monomeric MtrR engages a flexible loop within MtrA, stabilizing it into a short parallel β-sheet. In addition, MtrR interacts with the cytoplasmic domain of MtrA, on the face opposite the MtrA–MtrE interface and oriented toward the dimeric MtrH subunits. **A:** Overview of the Mtr-complex. Monomeric MtrR (orange) is predicted to interact with the MtrA (light blue) subunit of the complex. The hexamer of the MtrH subunit is depicted in green. Grey subunits represent MtrBCDEFG. **B:** PAE plot of the predicted aligned error for the complex prediction shown in A. The predicted interaction model between the Mtr complex and MtrR is of only moderate confidence, with further reductions observed when dimeric or trimeric forms of MtrR were included (data not shown). This likely reflects a broader limitation of AlphaFold3, which remains insufficiently trained for accurate structural prediction of small proteins and their interactions within larger complexes. **C:** The predicted model of the interaction between MtrA and MtrR colored by PLDDT **D:** The predicted model of the interaction between MtrR and MtrA shows MtrR interacting with the flexible MtrA-loop and the backside of the MtrA subunit, which is not interacting with MtrE or MtrH. Based on this prediction, MtrR might interact with the MtrA surface opposite of the cobalamin binding site and with the flexible loop of MtrA, which tethers the catalytic MtrA-domain to its membrane anchor. The MtrR binding to the flexible MtrA loop is potentially inhibiting the methyl-bound MtrA to interact with the MtrH, while further allowing the transfer to CoM. Consequently, these interactions are presumably not hindering the transfer of the methyl group in general, but might impact MtrA to favor a transfer of the methyl group onto CoM instead of H$_4$SPT. **E:** Detailed view of the interface of the MtrR-MtrA interaction. MtrR-24-28 are predicted to interact with MtrA-118 and MtrA-61-166. **F:** Schematic overview of the Mtr complex including MtrR modified after (Gottschalk & Thauer, 2001).

S1 Table: Used Strains and plasmids

| | | |
|---|---|---|
| **Archaea** | | |
| ***Methanosarcina mazei* Gö1** | Wildtype | German Collection of Microorganisms and Cell Cultures GmbH (DSMZ) number 3647 |
| ***Methanosarcina mazei* wt (3A)** | Wildtype,<br>improved growth on solid media | (Ehlers et al., 2005) |
| ***Methanosarcina mazei* ΔsORF16** | *M. mazei* 3A/ΔsP36 genomic deletion | This work |
| **Bacteria** | | |
| ***E.* coli DH5α** | host strain for cloning of plasmid constructs | (Hanahan, 1983) |
| ***E.* coli BL21 (DE3) containing pRIL** | strain for overexpression of genes with unusual codon usage; *Cm*$^R$ | Stratagene, La Jolla, USA |
| ***E. coli* C43 (DE3) containing pRIL** | strain for overexpression of hydrophobic or toxic proteins with unusual codon usage; *Cm*$^R$ | (Miroux & Walker, 1996) |
| **Plasmids** | | |
| **pmcl210** | cloning Vector, P15Aori, *lacZ*, *cat* | (Nakano et al., 1995) |
| **pRIL** | Encoding additional tRNAs | Stratagene, La Jolla, USA |
| **pET21a(+)** | expression vector, His-tag, *Amp*$^R$ | Novagen, Darmstadt, Germany |
| **pRS1736/alias pJS10** | expression vector, strep-tag, *Amp*$^R$ | Reiff-Trautmannsdorf et al., *in preparation* |
| **pRS207** | pac cassette in pBlueskript SK | (Ehlers et al., 2005) |
| **pRS1639** | *pMcrB;MMsORF16, Pur*$^R$*,* | (Tufail et al., 2024) |
| **pRS1743** | /MM_1547, *Pur*$^R$ | Reiff-Trautmannsdorf et al., *in preparation* |
| **pRS1834** | *pET21a/MMsORF16, Amp*$^R$ | This work |
| **pRS2037** | pWM321/MMsORF16, flanking regions, *Pur*$^R$ | This work |
| **pRS2087** | *Expression vector/ pMcrB, MMsORF16-His5, Pur*$^R$*,* | (Tufail et al., 2024) |
| **pRS2190** | *pRS1736/MM1543, Amp*$^R$ | This work |

**S2 Table: Used Oligonucleotides**

| construct | primer | Sequence (5’ to 3’) |
|---|---|---|
| ΔsORF16 | sORF16+1kb_for | TTTGCGGCCGCCTCCATCATAGTTGAGCTACCG |
| | sORF16+1kb_rev | TTTGGATCCGGAACAGCTGTCTCCCGC |
| | sORF16-1kb_for | TTTGGATCCGGCTCACCTTTGATTGAACTTTTG |
| | sORF16-1kb_rev | TTTGGTACCCTTTAAAATGGGGATTGCAGGTG |
| sORF16-$His_6$ | sORF16_nde_for | TTTCATATGAAAGAAATTATTACGCAAAAGCCTCTG |
| | sORF16_not1_rev | TTTGCGGCCGCGGTCTCCTCCGACTTTATG |
| strep-mtrA | mtrA_nde1_fwd | CATATGGCAGATAAAAGAGAACCAGC |
| | mtrA_not1_rev | GCGGCCGCTTACCTCCCTAGCAGTAATAACC |
